# Resolving Orsay Virus δ Protein Architecture Using Molecular Rulers in Single-Molecule Force Spectroscopy

**DOI:** 10.64898/2026.08.02.742377

**Authors:** Cynthia S. Deem, Sithara Wijeratne, Tsung-Cheng Lin, HanQiao Chen, Liaoqi Du, Yizhi Jane Tao, Ching-Hwa Kiang

**Affiliations:** Department of Physics & Astronomy, Rice University; Houston, Texas, 77005, USA; Department of Biosciences, Rice University; Houston, Texas, 77005, USA

## Abstract

Understanding the mechanical stability and architecture of viral proteins can provide valuable information about their biological function, but it remains a significant biophysical challenge. This study employs single-molecule force spectroscopy (SMFS) to investigate the multi-domain architecture of the Orsay virus δ protein, which lacks repeat structures and exhibits weak unfolding peaks. We engineered a construct using titin (I27)_4_ domains as an internal molecular ruler, enabling us to bracket the δ protein peaks to determine domain length and identify unfolding forces with an atomic force microscope (AFM). To address limitations of one-dimensional (1D) force distributions in resolving overlapping structural states, we created a two-dimensional (2D) mechano-structural signature map. By plotting kinetic stability (unfolding force *F*) against physical structural footprint (domain length *L*), we distinguished distinct unfolding domains, successfully separating degenerate 1D data into two statistically distinct populations corresponding to the δ protein’s internal domain (I) and C-terminal domain (C). This label-free method provides the first mechanical evidence of the δ protein’s multi-domain architecture. It establishes a robust, multi-dimensional framework for decoding the mechanics of complex biomolecular assemblies in their native state.

## Introduction

Single-molecule force spectroscopy (SMFS) is an effective technique[1–9] for investigating the mechanical and thermodynamic properties of various materials, including proteins[10–17], DNA[18–21], polymers[22], cells[23], and nanomaterials[24]. In SMFS experiments, a single molecule is anchored at both ends and stretched, allowing measurement of the spring’s restoring force. This process applies external forces that drive the system out of equilibrium, enabling direct observation of transitions between different states as the system adjusts to a new equilibrium. By analyzing the force peaks, researchers can gain valuable insights into the relative strength and size of domains that undergo forced unfolding[4,25]. This information is essential for identifying bond strengths within and between molecules, as well as for understanding the dynamics of protein folding.

Recent studies emphasize the role of SMFS in understanding viral protein mechanics, particularly at the SARS-CoV-2 spike-ACE2 interface[26], as well as in investigating cell adhesion[27]. Despite these successes, the application of SMFS to intact multidomain proteins in their native assemblies remains limited. Most studies have primarily focused on isolated single domains or repeats[10,28–30]. Investigating protein folding and the mechanics of a native multi-domain structure is a crucial next step. Isolating all domains can be challenging, and interactions between domains, as well as overall architecture, can significantly impact stability and function. These critical factors may be overlooked in studies focusing only on isolated domains.

The Orsay virus is the only naturally occurring viral pathogen of *C. elegans*, a small animal model widely used in research laboratories worldwide. Due to the ease of manipulation and the extensive resources available for studying *C. elegans*, the Orsay virus serves as an excellent model for investigating host-virus interactions. This virus features a novel fiber composed of a δ protein pentamer covalently linked to the capsid shell. The δ fiber plays a crucial role in binding to host receptors and facilitating viral entry[31]. At 420 Å in length, it consists of a fibrous shaft and two distinct globular domains: an internal domain (I) and a C-terminal domain (C) at the distal tip[32–34]. Understanding the mechanics and stability of these components is essential for comprehending their functions in the viral life cycle.

We used SMFS to study the full-length δ fiber of the Orsay virus in its native state, allowing us to examine its folding architecture and domain stability. To aid this process, we developed a fusion construct called δ-titin, where each δ protein subunit is flanked by four tandem titin I27 repeats (I27)_4_ at both the N- and C-termini. The titin I27 domain, which is the 27th Immunoglobulin-like (Ig) domain of the giant muscle protein titin, is now referred to as I91[35]. A construct made up of eight serially linked titin I27 domains, referred to as (I27)₈, has been used as a standard reference for force calibration in an atomic force microscope (AFM).

These titin I27 domains in the δ-titin construct serve as built-in nanoscopic rulers with well-characterized unfolding signatures, while also protecting the terminal ends of our target molecule when an AFM tip interacts with the molecule (Fig. 1). Similar to molecular tension sensors used to quantify force transmission in cellular environments[36], we used the titin I27 domains[1] as internal rulers to measure the unfolding mechanics of the multidomain δ fiber architecture. The use of tags on molecules for force spectroscopy has been explored[37,38]. By stretching individual δ-titin fusion constructs with AFM, we calibrated changes in domain length and identified unfolding events along the δ fiber.

**Fig 1.**
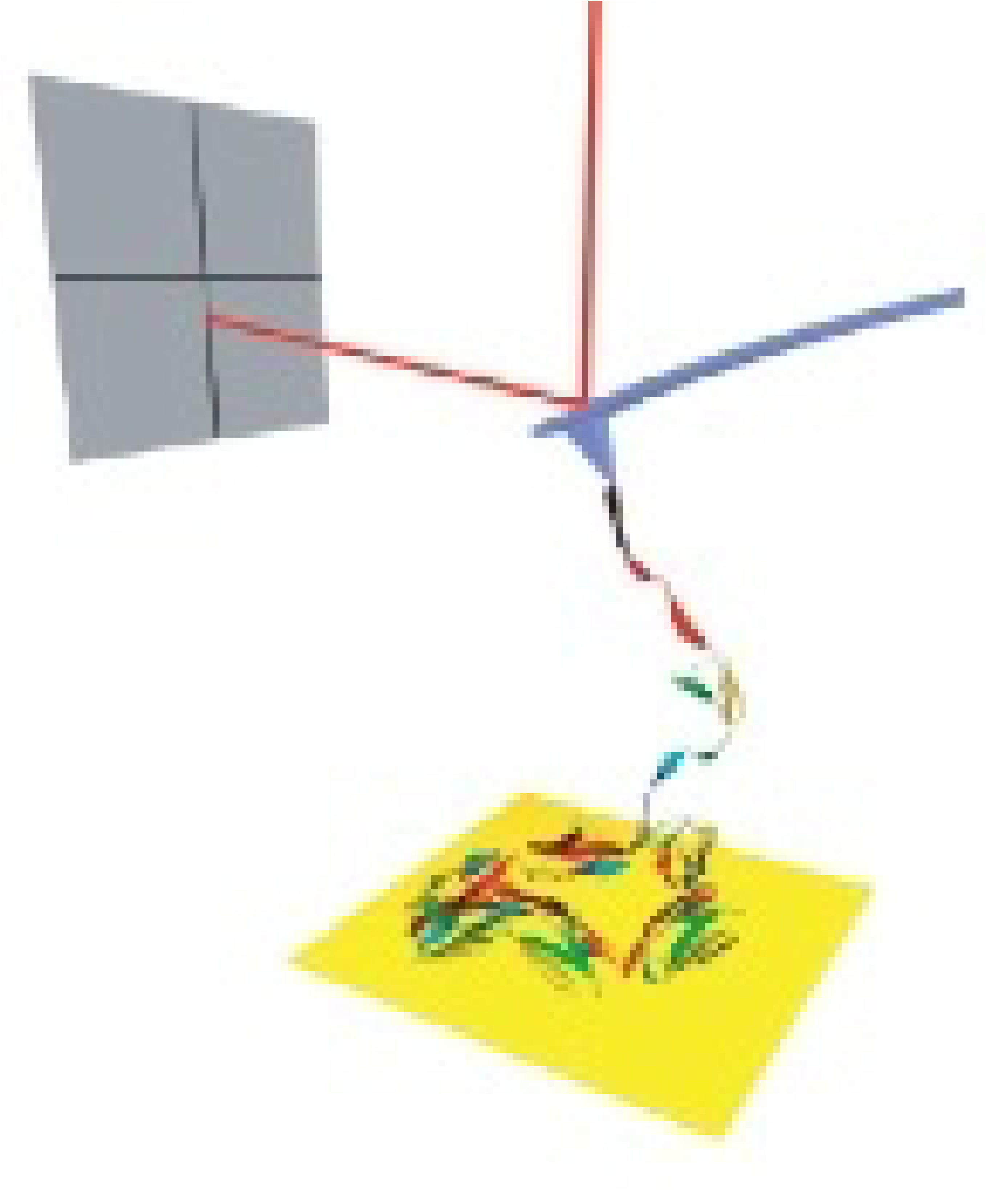
Illustration of AFM-based Single-Molecule Force Spectroscopy. The setup involves stretching a multidomain titin (I27)_8_ molecule. One part of the molecule binds to a cantilever tip, while the other connects to a gold substrate. As the piezo actuator moves the substrate away from the tip, the molecule is stretched. This label-free method can be applied to large multidomain proteins and was successfully used in this study on titin I27, δ fiber, and δ-titin, all while maintaining a consistent setup. The process stretches a single molecule, and the resulting force-distance curve reveals the unfolding events of the domains.

## Materials and Methods

### Bacterial culture conditions and selection

*E. coli* Rosetta competent cells were incubated on carbenicillin antibiotic plates at 37°C overnight. Afterward, a single colony was selected from the plate, transferred into 1/100 of LB medium containing 50 μg/ml carbenicillin, and shaken at 37 °C overnight.

### Construction of a recombinant strain

The genes corresponding to the full-length delta protein were amplified from Orsay viral DNA; the entire δ protein DNA segment was then cloned into the pEMI91 plasmid (74888, Addgene), which contains four titin I27 domains in both N-terminal and C-terminal, as well as the N-terminal His tag and C-terminal Strep tag, finally generating the His_6_-(I27)_4_-δ-(I27)_4_-Strep construction. The construct plasmid was transformed into *E. coli* Rosetta competent cells. PCR and DNA sequencing verified the construction strain.

### Protein expression and purification

To express proteins in *E. coli*, cells at the exponential growth phase were induced using Isopropyl β-D-1-thiogalactopyranoside (IPTG) to a final concentration of 0.4 mM when OD600nm reached 0.6-0.8. The protein was induced overnight at 16 °C and 220 rpm. The cell pellet was collected and then sonicated in a lysis buffer containing 50 mM Tris, pH 8.0, 300 mM NaCl, 10% (v/v) glycerol, 5 mM 2-Mercaptoethanol (2-ME), 1 mM NaN_3,_ then centrifuged at 30,000 xg for 45 minutes. The supernatant contained recombinant proteins purified using affinity chromatography on HisPur Ni-NTA resin (Thermo Scientific) and Strep-Tactin®XT resins (IBA Lifesciences). The eluted protein samples were injected into the Superose-6 (Cytiva) column. Afterward, the samples are run on an SDS-PAGE gel to determine which sample was purified most effectively. Purified protein fractions were subsequently concentrated using an Amicon® Ultra Centrifugal Filter (Millipore, UFC903008) to achieve a final concentration of 0.5 mg/mL.

### Western blot

The SDS-PAGE gel containing the pEMI91 δ protein was transferred to a polyvinylidene difluoride membrane, and the protein was analyzed by Western blot. The Western blot was performed with anti-strep primary antibody (1:5000, GT661, Invitrogen) and Goat anti-Mouse Secondary Antibody (1:5000, 31322, Invitrogen). First, the membrane was blocked with TBST containing 5% blocker for 60 minutes, then washed 3 times with TBST for 10 minutes each time. The membrane was then incubated with the primary antibody at a 1:5000 dilution for 2 hours. After washing, the membrane was incubated with the secondary antibody at a 1:5000 dilution for 40 minutes. After washing, the membrane was stained with NBT/BCIP substrate solution (34042, Thermo Scientific).

### Electron microscopy

For negative stain transmission electron microscopy (TEM), purified protein samples were diluted to 20–30 μg/mL and applied to Formvar/carbon 400-mesh copper grids (Cat# 01754-F, Ted Pella) that had been glow-discharged for 30 s at 15 mA using a PELCO easiGlow system (Ted Pella). The grids were stained with 7.5% uranyl formate for 45 s and blotted to remove excess stain. Imaging was performed on a JEOL JEM-1400 Flash transmission electron microscope equipped with a 15-megapixel AMT NanoSprint15 sCMOS camera. Micrographs were acquired at 30,000× magnification. Over 20 images were collected, with a representative micrograph shown.

### AFM experiments

The mechanical manipulation of titin (I27)_8_, δ, and the δ-titin fusion construct was conducted using a Nanoscope V Multimode VIII atomic force microscope (AFM) from Bruker. The protein samples were equilibrated at 37°C before being deposited onto a freshly prepared gold substrate, which was incubated at room temperature for 10 minutes. Any unbound material was removed by rinsing with phosphate-buffered saline (PBS, pH 7.4).

We used Bruker MLCT-O10 microlever probes, which are soft silicon nitride cantilevers and tips designed for high-load force-distance spectroscopy. Specifically, we selected the triangular C cantilever, with a nominal spring constant k = 0.01 N/m, a tip radius of 40 nm, a resonance frequency of 7 kHz, and dimensions of 310 μm × 20 μm. The precise spring constant was calibrated using the thermal noise method[39]. This technique involves measuring spontaneous, heat-induced Brownian vibrations while the cantilever is suspended above the surface. The AFM software then applies a Fourier transform to identify the resonance peak and calculates the spring constant using the Equipartition Theorem, which relates thermal motion to cantilever stiffness. To initiate single-molecule stretching, the AFM tip was brought into contact with the gold surface for 1 to 3 seconds, allowing non-specific protein adsorption. The typical surface contact time is 1 second, but we increased the duration when the local sample concentration is low; a longer contact time facilitates the capture of a complex.

Force measurements were conducted in PBS buffer containing 137 mM NaCl, 11.9 mM phosphate, and 2.7 mM KCl. These measurements were carried out at a pulling velocity of 1 μm/s. Protein immobilization was achieved through nonspecific binding rather than using covalent thiol-gold chemistry. This method avoids the need for chemical functionalization, which could compromise the native structural integrity of the viral fiber. While nonspecific binding can introduce variability in the exact orientation and pickup point, our engineered construct addresses this limitation. By sandwiching the δ protein between titin (I27)₄ handles, we ensure that the entire δ protein is likely to be captured within the stretched segment. Furthermore, in our experimental conditions, the nonspecific attachments are remarkably stable, enduring nanonewton-scale forces that exceed the internal unfolding forces of the targeted domains, thus facilitating single-molecule pulling. With a δ-titin sample concentration of 0.45 mg/ml, we maintained a low surface coverage, limiting the capture success rate to below 5%. This approach minimized the likelihood of pulling multiple molecules simultaneously. The observation of a single detachment peak in the force-distance curve confirmed that we were interacting with single molecules.

### Data Analysis and Polymer Mechanics Modeling

We initially analyzed the force-distance curves using a custom MATLAB program (MathWorks, Inc.) and then manually verified them. The inclusion criteria for identifying force peaks were that the force must be less than 300 pN, the domain lengths must be shorter than 30 nm, and the force-distance curves must exhibit at least three identifiable peaks. Only the ascending sections of the peaks that fit the WLC models were considered for identification.

We fit 1D histograms of *F* and *L* using both the Gaussian and stretched-exponential models[40], yielding similar peak values. To ensure mathematical consistency with the 2D analysis and to simplify the complexity of overlapping structural populations, we present the Gaussian distribution for clarity.

## Results

### Engineering a δ-titin Construct Using Titin (I27)_4_ Domains as Internal Molecular Rulers

Recombinant δ-titin protein (Fig. 2A), genetically engineered to contain four tandem I27 repeats at both the N- and C-termini, was expressed and purified by nickel-affinity and gel-filtration chromatography. Size-exclusion chromatography showed δ-titin eluted at a volume corresponding to a molecular mass over 670 kDa (Fig. 2B), consistent with the formation of a pentamer composed of 135 kDa δ-titin monomers of fibrous morphology. SDS-PAGE and Western blot analyses further confirmed the expected size and molecular identity of the purified fusion protein (Fig. 2C, left). Negative-stain transmission electron microscopy (TEM) revealed fibrous molecules with a prominent globular domain at one end and a smaller globular domain positioned along the fiber (Fig. 2D). This morphology closely matched that of the native δ fiber[41]. In addition, diffused shadows were observed at both ends, corresponding to the appended titin domains. Consistent with this interpretation, a histogram of the measured lengths of 138 δ-titin molecules showed an average length of 583 Å, significantly longer than 420 Å measured for the wild-type δ fiber (Fig. 2E)[41]. Together, these results indicate that δ-titin preserves the folded architecture and pentameric assembly of the wild-type δ fiber. The appended titin I27 domains do not significantly disrupt native δ fiber assembly, supporting the use of δ-titin as a suitable surrogate construct for single-molecule AFM pulling experiments.

**Fig 2.**
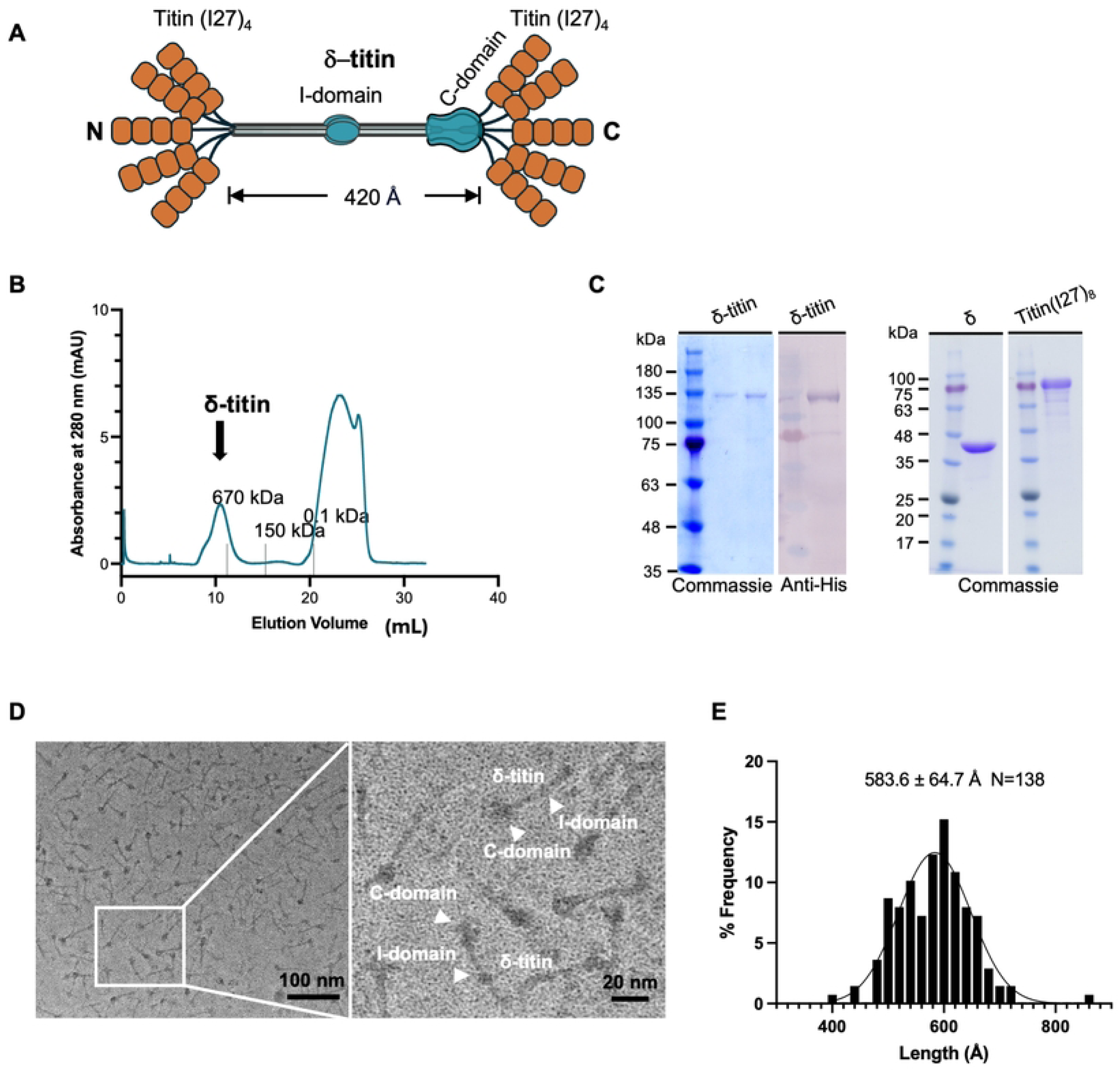
Construction and purification of the Orsay δ-titin protein. (A) Schematic of the δ-titin construct, with four titin I27 domains flanking each of the five full-length δ protein subunits at both N- and C-termini. (B) Gel filtration chromatogram of purified δ-titin using a Superose-6 column, with eluted positions of three molecular weight standards indicated. (C) SDS-PAGE analysis of purified δ-titin, δ, and titin (I27)_8_ used in this study. For δ-titin, both Coomassie-stained gels and western blot analyses are shown. (D) Negative-stain TEM of purified δ-titin with the C and I domains highlighted. (E) Histogram of δ-titin length measurements.

### Single-Molecule Force Spectroscopy (SMFS) using an Atomic Force Microscope (AFM)

We obtained force-distance curves for three sample types: a control polyprotein of eight titin I27 domains, the δ fiber alone, and the δ-titin fusion. Fig. 3A shows that the titin I27 octamer exhibited a sawtooth force curve with multiple distinct peaks. In contrast, the δ fiber alone produced only occasional low-force events with irregular spacings. These presumed δ unfolding peaks were weak and challenging to distinguish from background noise (Fig. 3B). The δ-titin fusion construct displayed a combination of both high-force and low-force peaks (Fig. 3C). This allowed unambiguous identification of δ unfolding events within the same titin I27 peaks, which function as internal calibration markers.

**Fig 3.**
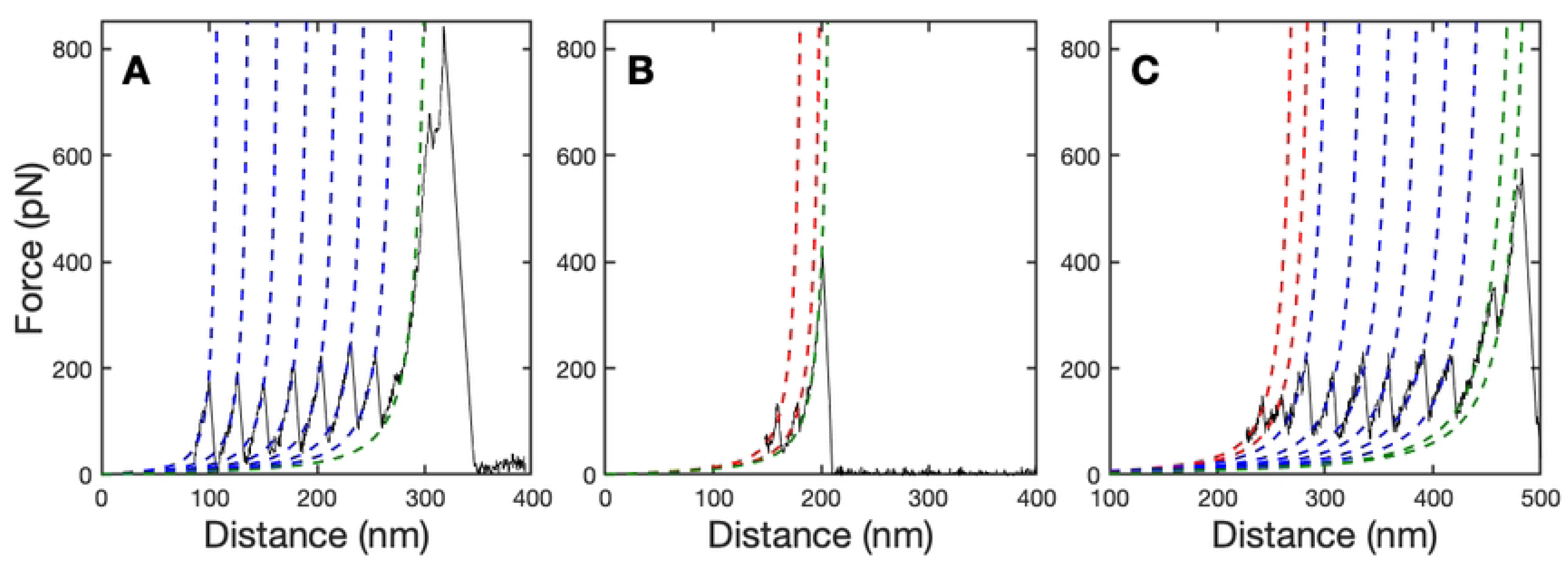
Representative single-molecule force-distance curves and worm-like chain (WLC) modeling. (A) The force-distance curve for the titin (I27)_8_ exhibits a characteristic sawtooth pattern. The WLC fits (blue dashed lines) confirm the sequential unfolding of individual titin I27 domains with regular contour-length increments. (B) The force-distance curve of the δ fiber (WLC fits shown as red dashed lines) displays smaller and irregularly spaced unfolding peaks that are difficult to determine. (C) The force-distance curve of the δ-titin fusion construct shows unfolding peaks corresponding to both the titin I27 domains and the δ domain. These force-distance curves illustrate the stretching of δ-titin pentamers. The well-characterized higher-force unfolding events of the titin I27 domains serve as an internal molecular ruler, successfully interleaving with those of the targeted δ domains. Solid lines represent experimental data, while dashed lines indicate the WLC fits to the unfolding curves. Red lines indicate δ domains, blue lines indicate titin I27 domains, and green lines indicate detachment peaks.

### Quantifying Domain Architecture via Worm-Like-Chain (WLC) Polymer Model

We utilize a label-free experimental setup that relies on nonspecific adsorption of molecules onto the cantilever and the gold substrate. This approach allows us to capture polyproteins, complexes, and fibrous assemblies without chemically functionalizing either the AFM tip or the target molecules. The mechanical attachment often demonstrates greater strength than DNA/RNA handles, allowing us to sustain forces in the nanonewton (nN) range, a regime typically unattainable with conventional methods that require chemical functionalization.

We use polymer mechanics models to analyze single-molecule force-distance trajectories, enabling the quantification of domain architecture dimensions. Before each unfolding transition, the cantilever restoring force exerted by the extended polypeptide was modeled using the WLC model[42],

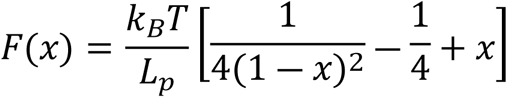

where *x* is the relative distance defined by *x = z/L_c,_ z* denotes the end-to-end distance, *L_c_* is the contour length, *F*(*x*) represents the cantilever restoring force at relative distance *x_,_ k_B_* is the Boltzmann constant, and *T* is the absolute temperature. We fixed the persistence length (*L_p_*) at 0.4 nm, reflecting the characteristic bending rigidity of unfolded polypeptides. By treating the contour length *L_c_* as a variable fitting parameter, we determined the molecule’s spatial dimensions across different mechanical states. The incremental change in contour length between successive force peaks at the peak position provides a direct experimental probe into the size of individual unfolded domains, which we define as the domain length (*L*). This metric, evaluated alongside the unfolding force (*F*), establishes the fundamental basis for our multi-dimensional mechanical analysis.

By using non-specific binding, we can resolve the native-state structure of the virus in its unmodified form, ensuring that the attachment remains stable under mechanical loads that are significantly greater than the internal unfolding forces of the domains. Because our experimental setup relies on highly stable, non-specific binding to study systems in their native state, the absolute AFM tip pickup point can occur at various locations along the polypeptide chain, leading to complex desorption signatures during the initial retraction phase. Because the relevant length scale is the relative distance, to reduce artifacts from nonspecific surface adhesion, the distance axis (*x* = 0) was aligned with the optimal offset parameter (*x_0_*) obtained from the initial WLC fit.

The necessity of the titin I27 internal ruler is further highlighted by our control experiments in which we stretch the untagged δ fiber. In the untagged setup, the C-terminal domain, δ(C), often serves as a nonspecific adhesion point to the substrate or the tip, leaving it outside the mechanically stretched segment. Consequently, its unfolding events are rarely observed. By flanking the δ with (I27)_4_ repeats, we effectively sequester the full δ(I) and δ(C) architecture within the active stretching segment. This engineering strategy dramatically enhances the probability of capturing the mechanical unfolding transitions of the native protein fiber.

We analyzed the distributions of *F* and *L* for each construct. Fig. 4 displays the one-dimensional (1D) histograms of peak *F* and the corresponding *L* obtained from repeated pulls. Titin I27-only control exhibited a narrow *F* distribution centered around 220 pN with a *L* of 25 nm per domain, which aligns with established values for titin I27. The data for δ alone showed a broad *F* distribution and an unresolved *L* distribution. The δ-titin fusion construct displays a bimodal distribution in both *F* and *L*.

**Fig 4.**
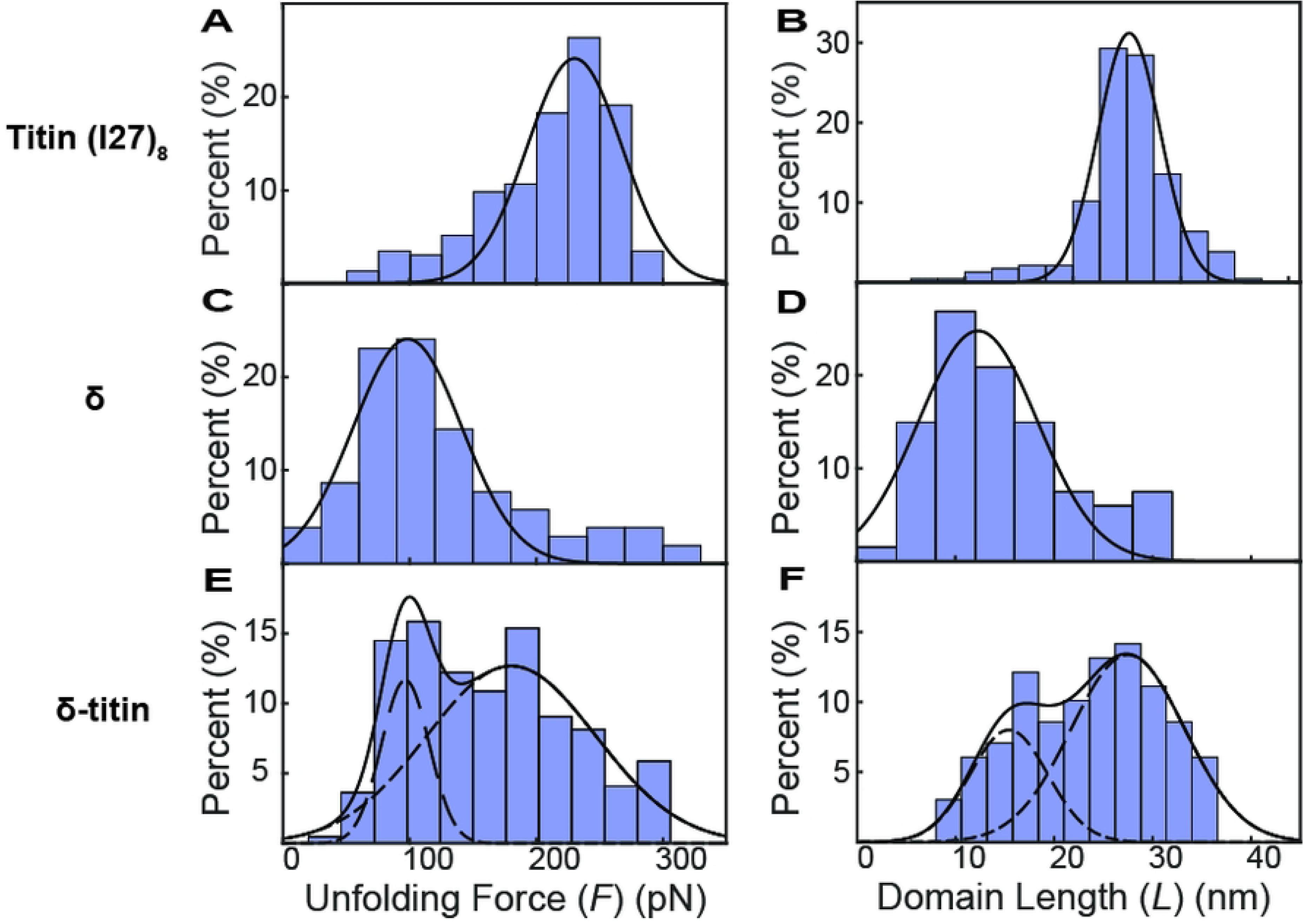
One-dimensional (1D) distributions of unfolding forces (*F*) and domain lengths (*L*). The distributions of titin (I27)_8_ show clear unimodal Gaussian populations in both (A) *F* and (B) *L*, consistent with previous studies (*n* = 236). In contrast, the δ fiber distributions show a broad, unresolved peak for both (C) *F* and (D) *L* (*n* = 59). The distributions for the δ-titin construct are bimodal in both (E) *F* and (F) *L* (*n* = 221); however, not all peaks are resolved (see text). All data were collected at a constant pulling velocity of 1 μm/s, and Gaussian distributions were used to identify distinct mechanical populations.

### Resolving Multi-Domain Architecture within a Two-Dimensional (2D) Mechano-Structural Signature Map

In standard SMFS analysis, evaluating kinetic stability solely using 1D *F* distributions often masks complex multi-domain architectures, as overlapping force regimes can obscure them. To overcome this limitation, we expanded our analysis to include a 2D mechano-structural signature map. In this map, the unfolding state of a specific protein domain is defined by two orthogonal parameters: the kinetic energy barrier required to unfold it, *F,* and its physical structural footprint, *L*. Fig. 5 presents a 2D contour map that illustrates the precise relationship between *F* and *L*. By representing the states orthogonally, this 2D visualization effectively converts the ambiguous 1D data into three distinct peaks.

**Fig 5.**
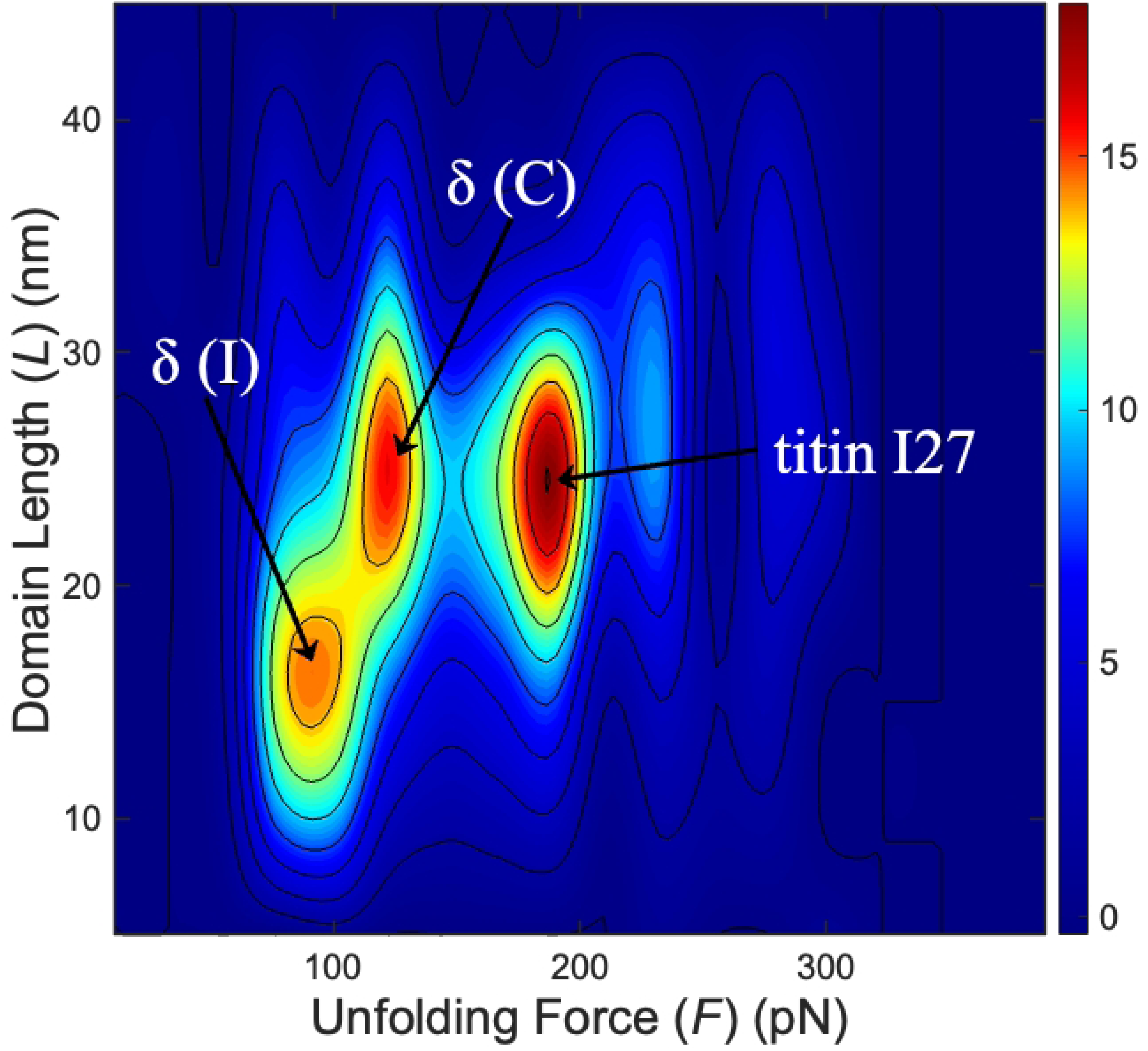
Two-dimensional (2D) contour plot of the unfolding force (*F*) versus domain length (*L*) of δ-titin. This 2D contour map resolves distinct mechanical populations that heavily overlap in standard 1D projections. The Gaussian Mixture Model (GMM) effectively distinguishes between the titin I27 domain, the C-terminal domain δ(C), and the internal domain δ(I). The (*F, L*) of titin I27, δ(C), and δ(I) are (210 pN, 24 nm), (130 pN, 29 nm), and (100 pN, 17 nm), respectively.

To identify peak locations, we applied a Gaussian Mixture Model (GMM) directly to the map. The GMM analysis confirmed the presence of a dense, highly stable titin I27 calibration cluster and clearly distinguished two adjacent, statistically distinct subpopulations within the lower-force regime. These mathematically resolved clusters correspond to the independent mechanical unfolding of the δ protein’s internal domain, δ(I), and its C-terminal domain, δ(C). This is consistent with structural studies that identified the δ protein as a pentameric fiber composed of an N-terminal coiled coil and a β-stranded fibrous shaft containing a small internal globular domain (I) and a larger C-terminal domain (C)[32,41]. Because coiled-coil structures typically unzip at relatively low forces of 8-15 pN[43], the N-terminal coiled-coil, and potentially portions of the β-stranded shaft, would unfold before the δ(I), δ(C), and titin I27 domains. Such low-force extension events would increase the initial contour length but would not generate distinct unfolding peaks in the force–distance curves; only the unfolding of the δ(I) and δ(C) domains would produce distinct unfolding peaks.

## Discussion

The measured domain length increments for the δ(I) and δ(C) domains were 17 nm and 29 nm, respectively. These values agree well with sequence-based predictions, indicating that the two domains contain approximately 41 and 106 residues, respectively[32], supporting the validity of our assignment strategy. By contrast, each I27 domain in δ-titin consists of 89 residues. Thus, the 2D mechano-structural signature map enables identification of the two structural domains within the δ fiber, which were otherwise difficult to distinguish using 1D length histograms alone.

The 2D SMFS characterization of the Orsay δ fiber clarifies its unique mechanical signatures, providing insight into the kinetic and thermodynamic properties of its constituent domains. By concurrently evaluating *F* and *L*, we resolved two unfolding populations that map to the anticipated δ(I) and δ(C) domains of the δ protein. Such results provide direct evidence for the multi-domain architecture of the δ protein, validate the existence of two discrete folding modules, and quantify their physical dimensions and stability in the native state. This versatile, label-free approach is broadly applicable to complex biomolecular assemblies, offering a critical mechanical perspective that complements structural insights derived from contemporary deep learning models.

This study highlights the effectiveness of SMFS, using titin-based molecular rulers, in characterizing protein domains in their native forms. By tagging the δ protein with titin (I27)_4_, we can distinguish between the δ(I) and δ(C) domains without isolating them or using repeated units. The two-parameter analysis that combines force and length provides a robust, generalizable framework for studying complex proteins. Recent advancements in artificial intelligence should enable a multiparameter approach to effectively identify and characterize multiple domains within complex structures. This innovative methodology opens new pathways for the study of complex systems. The unfolding forces observed are associated with transition state barriers, providing valuable insights into protein dynamics and enhancing our understanding of structure-function relationships.

This method illustrates the mechanical stability and domain architecture of the Orsay virus δ fiber. The unfolding forces of approximately 100 pN and 130 pN indicate moderate mechanical stability, suggesting that both the δ(C) and δ(I) domains are stably folded under force. This stability may be functionally important for the role of the δ fiber during Orsay virus infection. The δ(C) domain, which is hypothesized to mediate host receptor binding, likely presents a relatively rigid receptor-binding structure rather than undergoing major conformational rearrangements. Its mechanical stability may therefore help preserve the structural integrity of the receptor-binding region during attachment to the host cell surface. In contrast, the mechanical stability of the δ(I) domain may help maintain the δ protein as an extended, rigid fiber. This rigidity could allow the Orsay virus virion to project the C-terminal receptor-binding domain outward from the viral capsid and across the dense glycocalyx and collagen-containing extracellular matrix at the intestinal cell surface[44], thereby facilitating access to the putative host receptor. Thus, the measured unfolding forces support a model in which the δ fiber functions as a mechanically stable attachment structure, with the δ(I) domain contributing to fiber rigidity and the δ(C) domain mediating receptor engagement.

## Conclusions

In summary, this study sets a powerful precedent for examining a wide variety of proteins where domain interactions and stability are crucial to their function. From viral fibers to modular enzymes, this research improves our understanding of the stability of biological complexes and changes how researchers investigate protein folding and function.

## Acknowledgements

We thank the support from the E & M Foundation (CHK), NIH AI122356 (YJT), and the Welch Foundation C-1565 (YJT)

## Data availability

All data needed to evaluate the conclusions in the paper are present in the paper.

## Author contributions

CSD, SW, TCL, HC, and LD conducted experiments and analyzed the data; CHK and YJT designed the study; CHK, YJT, CSD, SW, and HC wrote the manuscript.

## References

1. Rief M, Gautel M, Oesterhelt F, Fernandez JM, Gaub HE. Reversible Unfolding of Individual Titin Immunoglobulin Domains by AFM. Science. 1997;276: 1109–1112. Available: https://www.jstor.org/stable/2893521

2. Kellermayer MSZ, Smith SB, Granzier HL, Bustamante C. Folding-Unfolding Transitions in Single Titin Molecules Characterized with Laser Tweezers. Science. 1997;276: 1112–1116. doi:10.1126/science.276.5315.1112

3. Oesterhelt F, Oesterhelt D, Pfeiffer M, Engel A, Gaub HE, Müller DJ. Unfolding Pathways of Individual Bacteriorhodopsins. Science. 2000;288: 143–146. doi:10.1126/science.288.5463.143

4. Harris NC, Song Y, Kiang C-H. Experimental Free Energy Surface Reconstruction from Single-Molecule Force Spectroscopy using Jarzynski’s Equality. Phys Rev Lett. 2007;99: 068101. doi:10.1103/PhysRevLett.99.068101

5. Bustamante CJ, Chemla YR, Liu S, Wang MD. Optical tweezers in single-molecule biophysics. Nat Rev Methods Primer. 2021;1: 25. doi:10.1038/s43586-021-00021-6

6. Chen W-H, Wilson JD, Wijeratne SS, Southmayd SA, Lin K-J, Kiang C-H. Principles of single-molecule manipulation and its application in biological physics. Int J Mod Phys B. 2012;26: 1230006. doi:10.1142/S021797921230006X

7. Walbrun A, Wang T, Matthies M, Šulc P, Simmel FC, Rief M. Single-molecule force spectroscopy of toehold-mediated strand displacement. Nat Commun. 2024;15: 7564. doi:10.1038/s41467-024-51813-9

8. Neupane K, Foster DAN, Dee DR, Yu H, Wang F, Woodside MT. Direct observation of transition paths during the folding of proteins and nucleic acids. Science. 2016;352: 239–242. doi:10.1126/science.aad0637

9. Barsegov V, Thirumalai D. Probing Protein-Protein Interactions by Dynamic Force Correlation Spectroscopy. Phys Rev Lett. 2005;95: 168302. doi:10.1103/PhysRevLett.95.168302

10. Wijeratne SS, Botello E, Yeh H-C, Zhou Z, Bergeron AL, Frey EW, et al. Mechanical Activation of a Multimeric Adhesive Protein Through Domain Conformational Change. Phys Rev Lett. 2013;110: 108102. doi:10.1103/PhysRevLett.110.108102

11. Wijeratne SS, Li J, Yeh H-C, Nolasco L, Zhou Z, Bergeron A, et al. Single-molecule force measurements of the polymerizing dimeric subunit of von Willebrand factor. Phys Rev E. 2016;93: 012410. doi:10.1103/PhysRevE.93.012410

12. Arce NA, Cao W, Brown AK, Legan ER, Wilson MS, Xu E-R, et al. Activation of von Willebrand factor via mechanical unfolding of its discontinuous autoinhibitory module. Nat Commun. 2021;12: 2360. doi:10.1038/s41467-021-22634-x

13. Serdiuk T, Manna M, Zhang C, Mari SA, Kulig W, Pluhackova K, et al. A cholesterol analog stabilizes the human β2-adrenergic receptor nonlinearly with temperature. Sci Signal. 2022;15: eabi7031. doi:10.1126/scisignal.abi7031

14. Yu H, Siewny MGW, Edwards DT, Sanders AW, Perkins TT. Hidden dynamics in the unfolding of individual bacteriorhodopsin proteins. Science. 2017;355: 945–950. doi:10.1126/science.aah7124

15. Solanki A, Neupane K, Woodside MT. Single-Molecule Force Spectroscopy of Rapidly Fluctuating, Marginally Stable Structures in the Intrinsically Disordered Protein α - Synuclein. Phys Rev Lett. 2014;112: 158103. doi:10.1103/PhysRevLett.112.158103

16. Choi H-K, Min D, Kang H, Shon MJ, Rah S-H, Kim HC, et al. Watching helical membrane proteins fold reveals a common N-to-C-terminal folding pathway. Science. 2019;366: 1150–1156. doi:10.1126/science.aaw8208

17. Takemasa, M., Sletmoen, M., and Stokke, B.T. Langmuir 25 (2009) 10174–10182.

18. Strunz T, Oroszlan K, Schäfer R, Güntherodt H-J. Dynamic force spectroscopy of single DNA molecules. Proc Natl Acad Sci. 1999;96: 11277–11282. doi:10.1073/pnas.96.20.11277

19. Smith DE, Tans SJ, Smith SB, Grimes S, Anderson DL, Bustamante C. The bacteriophage φ29 portal motor can package DNA against a large internal force. Nature. 2001;413: 748–752. doi:10.1038/35099581

20. Chen W, Chen W-H, Chen Z, Gooding AA, Lin K-J, Kiang C-H. Direct Observation of Multiple Pathways of Single-Stranded DNA Stretching. Phys Rev Lett. 2010;105: 218104. doi:10.1103/PhysRevLett.105.218104

21. Koch SJ, Wang MD. Dynamic Force Spectroscopy of Protein-DNA Interactions by Unzipping DNA. Phys Rev Lett. 2003;91: 028103. doi:10.1103/PhysRevLett.91.028103

22. Takemasa M, Sletmoen M, Stokke BT. Single Molecular Pair Interactions between Hydrophobically Modified Hydroxyethyl Cellulose and Amylose Determined by Dynamic Force Spectroscopy. Langmuir. 2009;25: 10174–10182. doi:10.1021/la9009515

23. Li J, Wijeratne SS, Nelson TE, Lin T-C, He X, Feng X, et al. Dependence of Membrane Tether Strength on Substrate Rigidity Probed by Single-Cell Force Spectroscopy. J Phys Chem Lett. 2020;11: 4173–4178. doi:10.1021/acs.jpclett.0c00730

24. Wijeratne SS, Penev ES, Lu W, Li J, Duque AL, Yakobson BI, et al. Detecting the Biopolymer Behavior of Graphene Nanoribbons in Aqueous Solution. Sci Rep. 2016;6: 31174. doi:10.1038/srep31174

25. Jarzynski C. Nonequilibrium Equality for Free Energy Differences. Phys Rev Lett. 1997;78: 2690–2693. doi:10.1103/PhysRevLett.78.2690

26. Bauer MS, Gruber S, Hausch A, Melo MCR, Gomes PSFC, Nicolaus T, et al. Single-molecule force stability of the SARS-CoV-2–ACE2 interface in variants-of-concern. Nat Nanotechnol. 2024;19: 399–405. doi:10.1038/s41565-023-01536-7

27. Jo MH, Meneses P, Yang O, Carcamo CC, Pangeni S, Ha T. Determination of single-molecule loading rate during mechanotransduction in cell adhesion. Science. 2024;383: 1374–1379. doi:10.1126/science.adk6921

28. Cecconi C, Shank EA, Bustamante C, Marqusee S. Direct Observation of the Three-State Folding of a Single Protein Molecule. Science. 2005;309: 2057–2060. doi:10.1126/science.1116702

29. Staple DB, Payne SH, Reddin ALC, Kreuzer HJ. Model for Stretching and Unfolding the Giant Multidomain Muscle Protein Using Single-Molecule Force Spectroscopy. Phys Rev Lett. 2008;101: 248301. doi:10.1103/PhysRevLett.101.248301

30. Zocher M, Bippes CA, Zhang C, Müller DJ. Single-molecule force spectroscopy of G-protein-coupled receptors. Chem Soc Rev. 2013;42: 7801–7815. doi:10.1039/C3CS60085H

31. Félix M-A, Ashe A, Piffaretti J, Wu G, Nuez I, Bélicard T, et al. Natural and Experimental Infection of Caenorhabditis Nematodes by Novel Viruses Related to Nodaviruses. PLOS Biol. 2011;9: e1000586. doi:10.1371/journal.pbio.1000586

32. Guo YR, Fan Y, Zhou Y, Jin M, Zhang JL, Jiang H, et al. Orsay Virus CP-δ Adopts a Novel β-Bracelet Structural Fold and Incorporates into Virions as a Head Fiber. J Virol. 2020;94: doi:10.1128/jvi.01560-20

33. Franz CJ, Renshaw H, Frezal L, Jiang Y, Félix M-A, Wang D. Orsay, Santeuil and Le Blanc viruses primarily infect intestinal cells in Caenorhabditis nematodes. Virology. 2014;448: 255–264. doi:10.1016/j.virol.2013.09.024

34. Guo YR, Hryc CF, Jakana J, Jiang H, Wang D, Chiu W, et al. Crystal structure of a nematode-infecting virus. Proc Natl Acad Sci U S A. 2014;111: 12781–12786. doi:10.1073/pnas.1407122111

35. Bang ML, Centner T, Fornoff F, Geach AJ, Gotthardt M, McNabb M, et al. The complete gene sequence of titin, expression of an unusual approximately 700-kDa titin isoform, and its interaction with obscurin identify a novel Z-line to I-band linking system. Circ Res. 2001;89: 1065–1072. doi:10.1161/hh2301.100981

36. Wang X, Ha T. Defining Single Molecular Forces Required to Activate Integrin and Notch Signaling. Science. 2013;340: 991–994. doi:10.1126/science.1231041

37. Giganti D, Yan K, Badilla CL, Fernandez JM, Alegre-Cebollada J. Disulfide isomerization reactions in titin immunoglobulin domains enable a mode of protein elasticity. Nat Commun. 2018;9: 185. doi:10.1038/s41467-017-02528-7

38. Sun Y, Liu X, Huang W, Le S, Yan J. Structural domain in the Titin N2B-us region binds to FHL2 in a force-activation dependent manner. Nat Commun. 2024;15: 4496. doi:10.1038/s41467-024-48828-7

39. Hutter JL, Bechhoefer J. Calibration of atomic-force microscope tips. Rev Sci Instrum. 1993;64: 1868–1873. doi:10.1063/1.1143970

40. Dudko OK, Hummer G, Szabo A. Theory, analysis, and interpretation of single-molecule force spectroscopy experiments. Proc Natl Acad Sci U S A. 2008;105: 15755–15760. doi:10.1073/pnas.0806085105

41. Fan Y, Guo YR, Yuan W, Zhou Y, Holt MV, Wang T, et al. Structure of a pentameric virion-associated fiber with a potential role in Orsay virus entry to host cells. PLOS Pathog. 2017;13: e1006231. doi:10.1371/journal.ppat.1006231

42. Marko JF, Siggia ED. Stretching DNA. Macromolecules. 1995;28: 8759–8770. doi:10.1021/ma00130a008

43. Bornschlögl T, Rief M. Single Molecule Unzipping of Coiled Coils: Sequence Resolved Stability Profiles. Phys Rev Lett. 2006;96: 118102. doi:10.1103/PhysRevLett.96.118102

44. Zhou Y, Chen H, Zhong W, Tao YJ. Collagen and actin network mediate antiviral immunity against Orsay virus in C. elegans intestinal cells. PLOS Pathog. 2024;20: e1011366. doi:10.1371/journal.ppat.1011366

